# Reconstructing Great Basin Butterfly-Pollen Interaction Networks over the Past Century

**DOI:** 10.1101/2020.12.25.424352

**Authors:** Behnaz Balmaki, Tara Christensen, Lee A. Dyer

## Abstract

**Aims:** Insects and the plants they interact with dominate terrestrial biomes and constitute over half of the earth’s macro-organismal diversity. Their abundance in museum collections can provide a wealth of natural history data if they are collected as part of careful ecological studies or conservation programs. Here, we summarize pollen-insect quantitative networks gleaned from adult lepidopteran museum specimens to characterize these interactions and to examine how richness and frequency of butterfly-pollen associations have changed over a 100-year time series in Nevada and California. Pollen collected from well-curated butterfly specimens can provide insight into spatial and temporal variation in pollen-butterfly interactions and provide a complement to other approaches to studying pollination, such as pollinator observation networks.

**Location:** Great Basin and Sierra Nevada: California, Nevada

**Time period:** The last 100 years

**Major Taxon studied:** Butterflies

**Methods:** We estimated butterfly-pollen network parameters based on pollen collected from butterfly specimens from the Great Basin and Sierra Nevada. Additionally, we pooled interaction networks associated with specimens captured before and after 2000 to compare pollen-pollinator interaction variation under drought periods in California and Nevada in the last two decades versus previous years in the time series.

**Results:** Butterfly-pollen networks indicated that most pollen-butterfly species interactions are specialized and appear to be different from observational networks. Interaction networks associated with specimens captured before and after 2000 revealed that compared to previous decades, butterfly-pollen networks over the past 20 years had higher nestedness and connectance, with high pollen richness and low pollen abundance.

**Main conclusions:** These findings represent another unique approach to understanding more about pollination biology, and how butterfly-pollen interactions are impacted by climate variation and ecosystem alteration.

## Introduction

Plant-pollinator interactions contribute substantially to biodiversity and are likely to provide ecosystem stability in the face of climate change and environmental stress. Documenting interactions between plants and pollinators is the first step in resolving different aspects of pollinators’ habitat, diet, migration, and impact on plant reproduction (Jones, 2012, 2014). Pollination interaction networks are an increasingly important method for summarizing these interactions, and they are frequently used to document and describe plant-visitor interactions, pollen-insect interactions, and pollinator effectiveness (Ballentyne et al., 2015; Tur et al., 2016). Numerous metrics at both the network and species level have been developed and can provide useful quantitative summaries of pollination network properties (Gibson et al., 2011; Jauker, et al., 2018). Proper interpretations of selected network parameters can help us to understand pollinators’ responses to habitat alteration, biodiversity change, and climate change by comparing changes in interactions over time or across disturbance gradients (Proulx et al., 2005; Ballantyne et al., 2015; Bohan & Dumbrell, 2017). Analyzing network indices is also a useful tool for assessing the level of generalization or specialization for species on either side of the bipartite network. For example, published pollination networks have been used to estimate diet breadths of pollinators that are on a continuum ranging from extreme specialists (one link between the pollinator and its host plant), to extreme generalists (many links between the pollinator and its host plants), but the estimates are based solely on network structure (Dormann et al., 2009; Bohan & Dumbrell, 2017; Novella-Fernandez et al., 2019). These networks are usually “visitation networks,” utilizing observations of insects visiting flowers, rather than analysis of pollen collected from insects.

Pollen analysis is commonly used in the reconstruction of past climate, vegetation history, historical ecology, and biodiversity (Matthias et al., 2015; Shennan et al., 2015; Balmaki et al., 2019). Palynological methods allow for comparisons of past and modern plant diversity and species assemblages (Gosling et al., 2018), providing quantitative measures of landscape change through time that can be linked to landscape disturbance or other perturbations to investigate changes in plant communities with a focus on the impacts of climate change on natural and managed ecosystems (Burjachs & Julia, 1994; Balmaki et al., 2019; Díaz et al., 2019). Analysis of pollen associated with insects is a more recently developed method that can be used for addressing questions related to insect food sources and habitats (Jones, 2012; Silberbauer et al., 2004), but this approach has not been utilized for examining butterfly-pollen networks. Pollen persists on lepidopteran bodies through collection and curation (Courtney et al., 1982), so analyzing pollen grains on butterflies can provide information on insect-pollen interactions that is distinct from existing pollination network approaches. Historically, flower visitation networks have served as the most common method for examining plant-pollinator interactions; for these networks, visitors to flowers of a particular plant are recorded and interaction frequency is used as a proxy for pollination. However, visitation does not always correspond to successful pollen transfer, and this method can be an ineffective means of predicting the importance of particular pollinators (King et al., 2013). Some visitors do not pick up pollen, simply remove pollen without transferring it to another conspecific, or interfere with pollination by blocking the stigma of the flower (Ballantyne et al., 2015). More advanced methods of studying plant-pollinator interactions have emerged in recent years, and present more precise alternatives to studying flower visitation (Garratt & Potts, 2011; O’Connor et al., 2019).

One of the most precise methods for documenting interactions between insects and pollen (i.e. not pollination per se) is to merge identification methods from palynology with samples collected from insects that have been identified and curated. Museum specimens provide permanent records of these interactions (Colla & Packer., 2008; Colla et al., 2012; Bartomeus et al., 2013; Scheper et al., 2014). Upon nectaring at flowers, butterflies make contact with the reproductive organs of flowers and pollen is lodged onto their bodies. While butterflies are foraging for nectar with their elastic proboscis, pollen grains stick to their eyes, proboscis, frons, antennae, wings, and legs, which renders them as pollen carriers and potential pollinators (Willmer, 2011). Identification and quantification of the pollen grains found on the bodies of butterflies or other pollinators can provide information about the plants that an individual has come into contact with and can be used to create plant-pollen networks for various inferences, including assumptions that these networks estimate pollination networks (Butler & Johnson, 2020). Although the presence of pollen on insects is an imperfect indicator of which plants the insect provides pollination services to, it suggests an interaction with the reproductive floral parts of that species (Jennersten, 1984; Butler & Johnson, 2020). This method may provide an insightful alternative to the visitation networks that dominate the pollination network literature (Silberbauer et al., 2004; Kleijn & Raemakers, 2008; Jones, 2012; Scheper et al., 2014) since insects in visitation networks can include parasites and incidental visitors that are not picking up or transferring pollen (Ballantyne et al., 2015). Examining pollen on preserved insects is also a step towards characterizing insect communities that are carrying and moving pollen at any given place and time.

Natural history museums are an important tool for recording taxonomic diversity and can provide insight into native pollinator communities across spatial and temporal gradients (Titeux et al., 2017; Seltmann et al., 2017). Museums and their collections are also a key part of efforts to document and predict the consequences of habitat loss, fragmentation, invasive species, and climate change. The few studies that have used museum specimens to quantify insect movements and temporal changes in pollinators have mostly focused on Hymenoptera (Silberbauer et al., 2004; Wood et al., 2019) and have effectively documented striking spatial and temporal patterns. For example, Wood et al. (2019) found that specimens from declining bumblebee species exhibited a one-third decrease in pollen richness from individuals collected before and after 2000. Additional studies using museum specimens from different systems and with diverse pollinator taxa will contribute to understanding how plant-pollen interactions and pollination networks are responding to global change.

Overall, weather patterns in the Sierra Nevada and Great Basin are rapidly changing, and these changes have been especially severe in the last two decades (McEvoy et al., 2012; Hatchett et al., 2015). Substantial changes include the decreasing snowpack, decreases in annual precipitation, and increased fire frequency (Belmecheri et al., 2016). Concurrently, there have been considerable increases of invasions by exotic plants that are outcompeting the endemic species over the past 100 years. The proliferation of drought resistant and fire tolerant plants has contributed to a decline in the density of native plants which are not adapted to these climate extremes (Rondeau, 2013). These changes in plant composition necessarily result in shifting pollinator communities that interact with them. One of the primary aims of this study was to utilize plant-insect interaction data associated with the Lepidoptera collections at the University of Nevada Museum of Natural History (UNRMNH) to examine how richness and abundance of butterfly-pollen interactions have changed in the Great Basin and Eastern Sierra over the last century. This time period and sampling area include multiple disturbance gradients, but the focus here was on pre and post 2000 pollination networks. In order to investigate the effects of time on the structure and composition of pollination networks in California and Nevada, we used specimens from the invertebrate collections at the University of Nevada, Reno Museum of Natural History. These specimens were all adults from the Lepidopteran superfamily, Papilionoidea, commonly known as butterflies (Kawahara et al., 2019), which were collected between the years of 1910 and 2020. Pollen grains were removed from these specimens and identified to the species or genus level. We quantified the pollen counts and calculated commonly used network parameters based on the species interaction matrix. To investigate pollen-pollinator interaction variation under drought conditions in California and Nevada in the last two decades, networks before and after 2000 were used to examine changes in network structure over time.

## Methods

### Data Collection and Pollen Analysis

One hundred seventy-three Lepidopteran specimens from the most common species sampled over the past century were selected from the UNR Museum of Natural History (UNRMNH) to collect pollen grains from their bodies. These specimens were collected between 1910-2020 in the Great Basin region of Nevada as well as the Sierra Nevada mountain range in Nevada and California (Table S1). The samples represented three papilionoid (butterfly) families in Lepidoptera and included 18 different species. Specimens were rinsed with 95% ethyl alcohol (ETOH) to collect all pollen grains from their external tissues, including the proboscis, legs, and compound eyes (Fig.1). Entomological pins were used to remove all the pollen grains from each insect’s external tissue under a binocular microscope. All recovered pollen grains were stored in vials with 2000 cs silicone oil volume and stain (Safranin-O) to highlight morphological features (Jones, 2014). Representative pollen samples were mounted on glass slides by adding two drops of the sample solutions to a clean glass slide, securing with a coverslip, and sealing with clear nail polish. Light microscopy (LM) and a scanning electron microscope (SEM) were used for pollen identification. In addition, the Great Basin pollen database at the UNRMNH was used as a reference slide for pollen identification.

**Figure 1.**
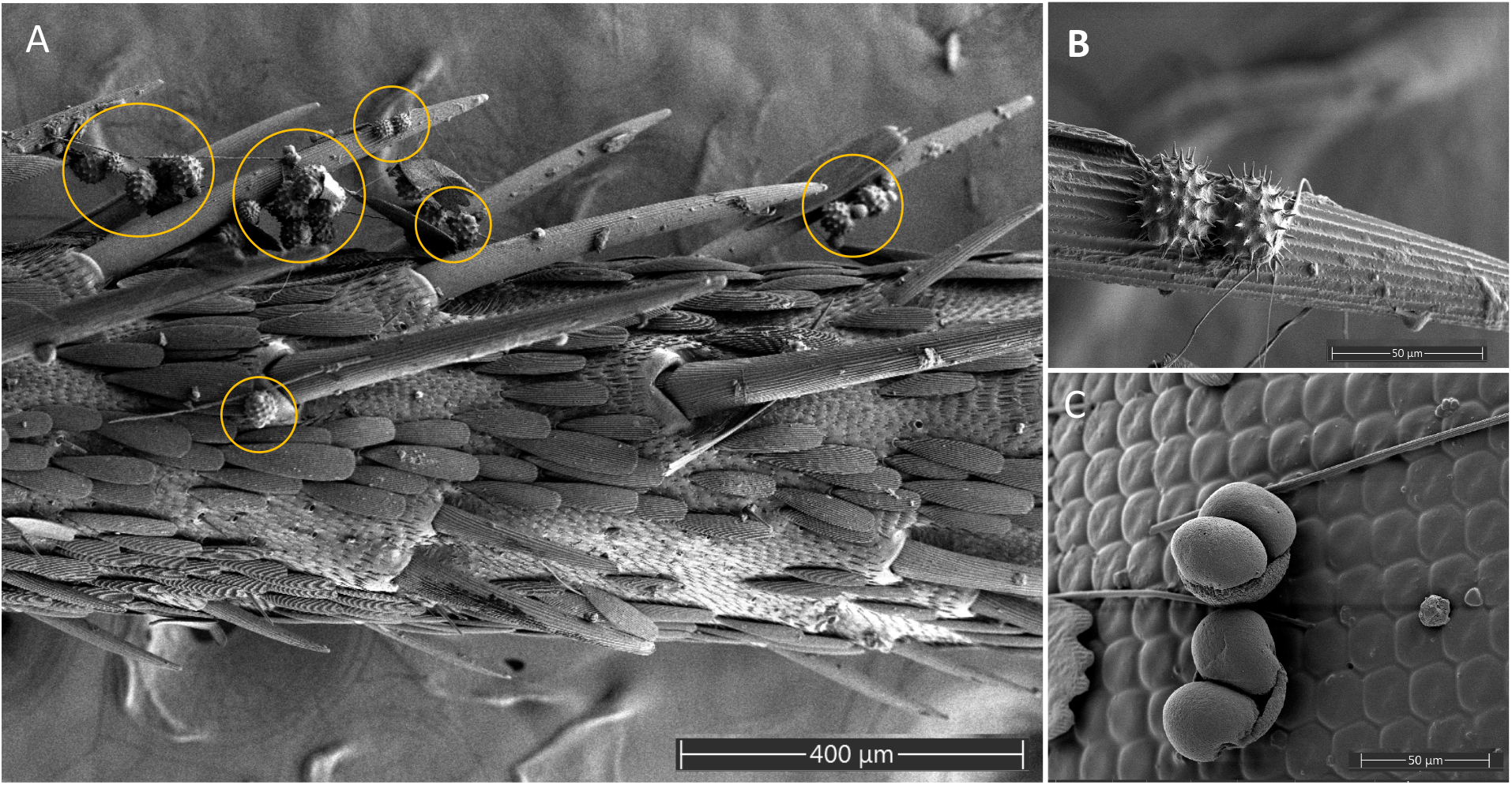
A-C: Scanning electron micrograph of pollen grains on butterfly legs and eyes from the collection at the UNRNHM. A-B: Asteraceae pollen covering the leg of a nymphalid butterfly. C: Pine pollen on a butterfly eye.

### Network and Statistical Methods

For each specimen, we quantified pollen species abundances and unique interaction frequencies for each available year from 1910 to 2020 to compare patterns in diversity and abundance across time. Well-established protocols were used for visualizing ecological networks and calculating network parameters (e.g., Pardikes et al., 2018; Dell et al., 2019; Salcido et al., 2020), and networks were analyzed using the “bipartite” package in R (Fig. 2, Appendix 1 Figure A1). The nodes in the bipartite network represent the plant and pollinator (i.e. pollen-carrying) species, and the edges were determined by pollen-insect associations present in museum specimens, with the width of edges representing the frequency of encounter of a given interaction. We calculated network indices including connectance, nestedness, network specialization (H2’), as well as other commonly reported network metrics (Table S2). Connectance represents the number of links between nodes over the number of species squared in a network; this parameter summarizes the number of realized possible connections (Martinez, 1992). Nestedness describes the degree of subsetting; compared to random networks, nested networks contain more specialized interactions as distinct subsets of more generalized interactions. Different parameters have been developed for nestedness, for example the NODF parameter, for which a value of 100 represents a perfectly nested network, and a value of 0 represents no nestedness. Network specialization is a network-level index that summarizes the degree of specialization and is useful for comparisons across multiple networks. Values of network specialization range between zero and one, with zero representing complete generalization and one representing complete specialization (Blüthgen 2006).

**Figure 2.**
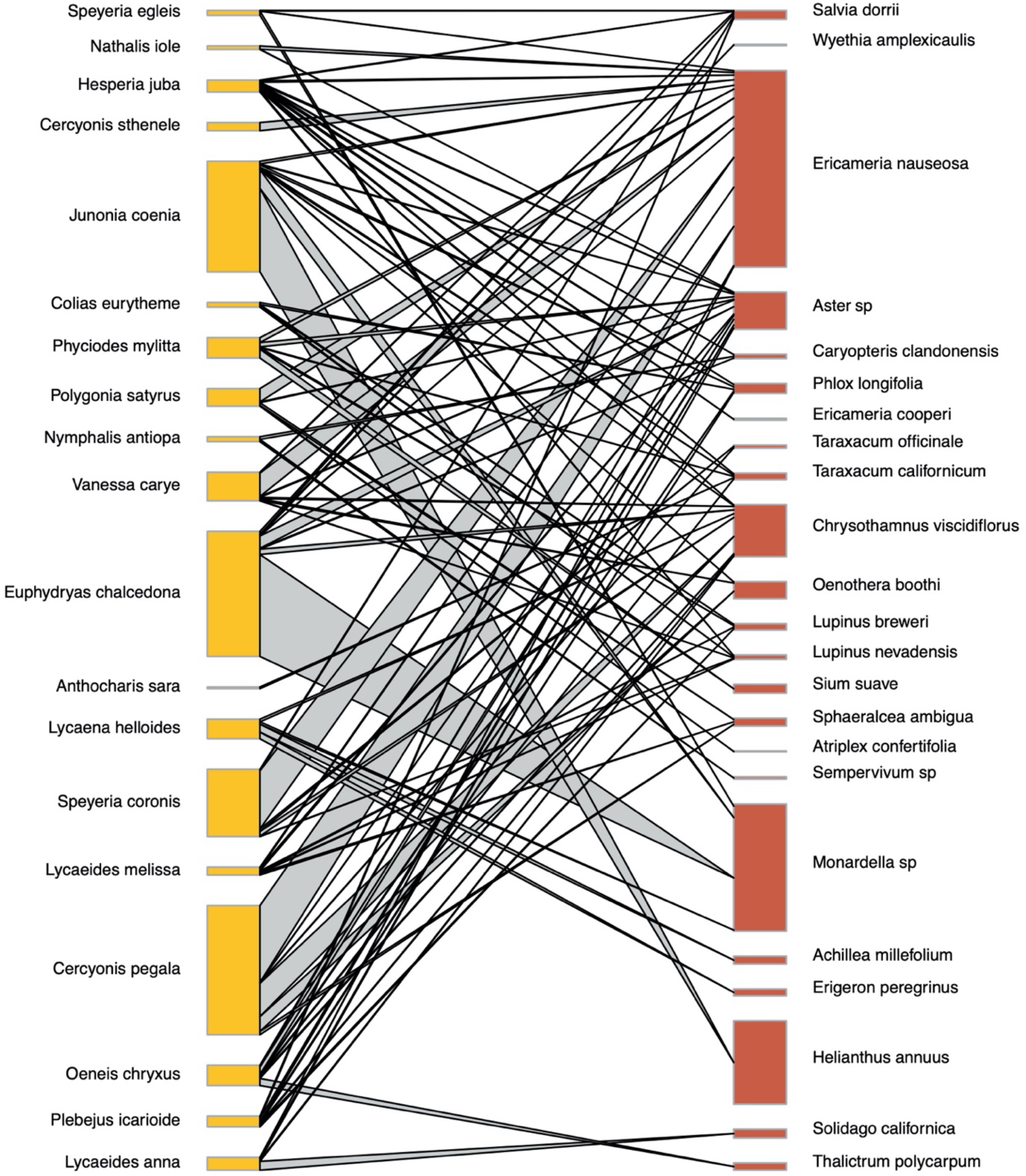
Bipartite pollination network of 19 pollinators and pollen from 29 plant species. Links between plants and pollinators are represented with lines whose width is proportional to the number of interactions while the width of the nodes represents total abundance of that taxon across all of its interactions.

Based on warming events which impacted California and Nevada around 2000 (McEvoy et al., 2012; Hatchett et al., 2015; Belmecheri et al., 2016), we considered two distinct pollen-pollinator interaction networks; one using the data collected before 2000, and one using the data collected after 2000 (Fig. 3). We calculated network parameters for these two groups to compare the level of specialization for the selected butterflies over time considering the changing climate in these two time periods (Table 1). Also, the richness values were included as response variables in Bayesian linear models to examine changes in diversity over time for individual species (details are in the supplementary materials, Appendix 2 Figure A2). Networks were not compared to null networks due to the descriptive nature of the study. To improve the relevance of results to pollination, plant species known to be wind-pollinated were removed from the analysis, since it is likely insects picked up their pollen in the environment rather than making contact with them.

**Figure 3.**
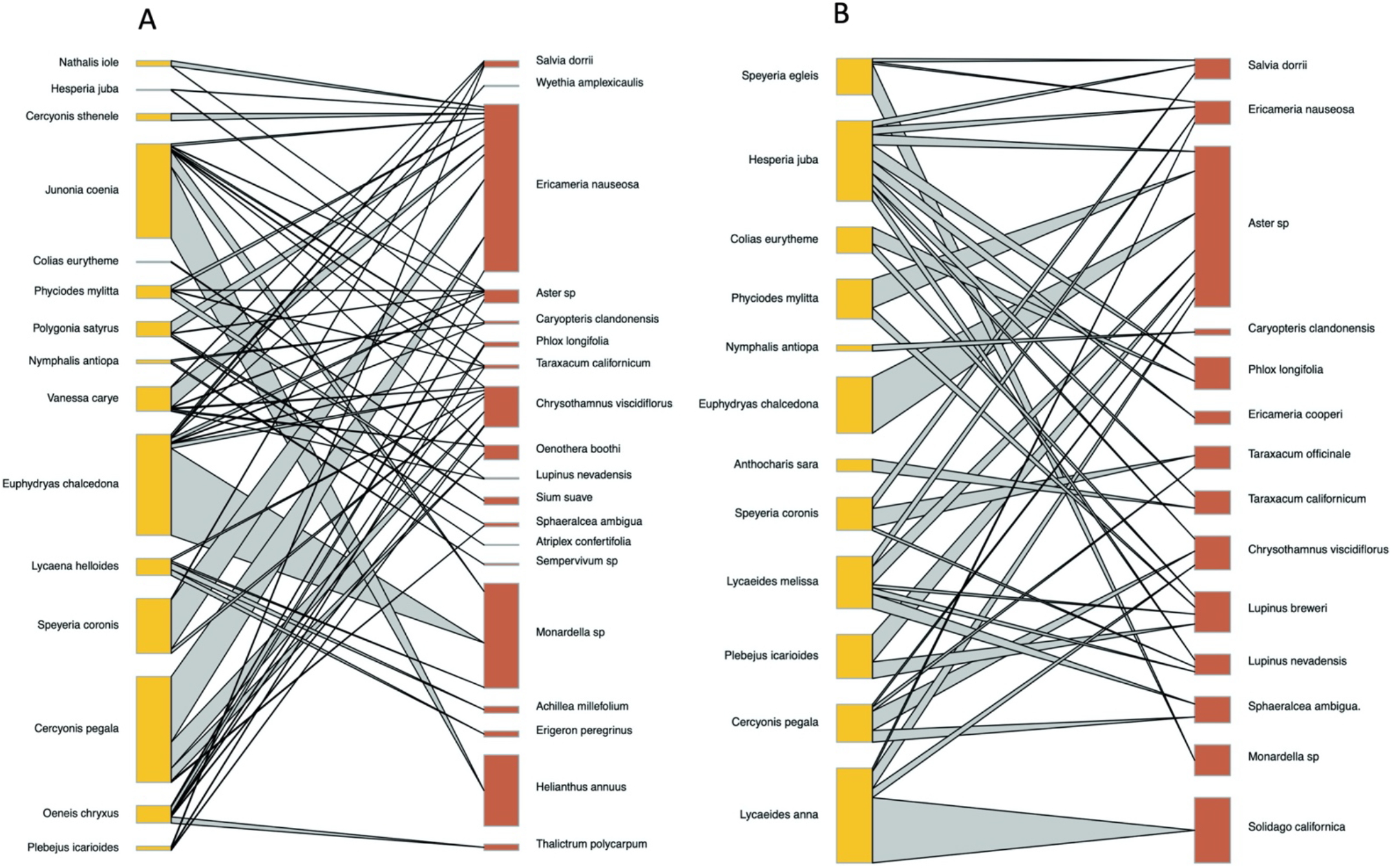
Bipartite pollination networks A: Before 2000, B: After 2000.

**Table 1.**
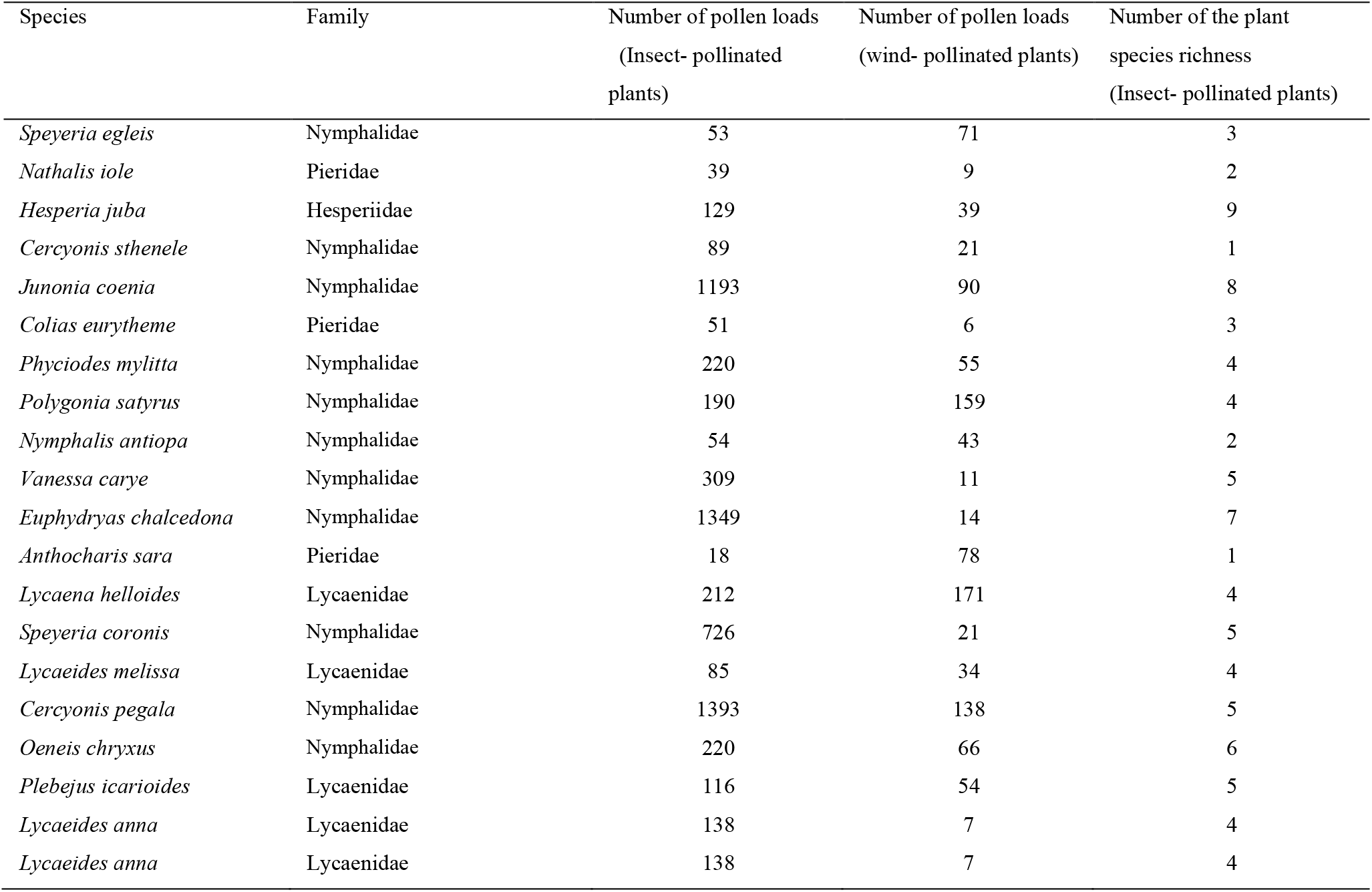
list of the butterflies species, number of pollen loads and number of plant species richness on each buttefly species.

### Pollinator observations

In order to better understand plant-pollinator interactions for some of the species examined here, and to provide a brief qualitative comparison to the pollen counts found on individuals, short targeted pollinator observations were conducted and combined with existing observational knowledge. Two species, *Lycaeides anna* and *L. melissa*, were selected due to our familiarity with the natural history of these species as well as their high abundances at the selected sites. A population of *L. anna* was observed near Yuba Pass in eastern California, and a population of *L. melissa* was observed in Verdi, Nevada on August 3, 2020 and August 11, 2020, respectively. The host plant for *L. anna* larvae at this site is *Lotus nevadensis* and for *L. melissa* the host plant is *Medicago sativa*. At each site, the perimeter of the sampling area was walked continuously for two hours. Each time an individual was encountered, the sex of the individual was recorded, and it was noted whether the individual was found on the host plant or not. In the case of *L. anna*, the identities for all other plants visited were recorded, and all individuals found on plants were observed visiting the flowers of those plants. Due to the great abundance and activity of *L. melissa* at the other site, it was only possible to record whether individuals landed on their host plant or were observed flying or on another plant, and individuals were found visiting flowers as well as landing on stems or leaves. After the two-hour observation window, individuals were collected opportunistically for pollen analysis. A total of 13 individuals from *L. anna* and 20 individuals from *L. melissa* were captured and frozen for pollen analysis, which was completed using the same methods described for the network assessment (Appendix 3 Figure A3).

## Results

### Pollen Analysis of Historic Lepidoptera

We identified a total of 7686 individual pollen grains on the external tissues of our butterfly specimens. Most of the pollen was found on the eyes, proboscis, and legs of the butterflies, and we accounted for pollen from 29 different plant species. Pollen analysis from as far back as 1910 to the present revealed a diverse community of plant species commonly associated with butterflies (Table 1). Pollen from *Ambrosia dumosa* (Asteraceae)*, Artemisia tridentate* (Asteraceae)*, Pinus* sp (Pinaceae)*, Atriplex confertifolia* (Amaranthaceae)*, Alnus tenuifolia* (Betulaceae) and *Leymus cinereus* (Poaceae) were found mostly on the butterflies’ head and wings. These species are not known to be insect pollinated, and it is likely these insects inadvertently picked up their pollen from the environment; perhaps in the soil or from the wind (Blauer et al., 1976). Overall, our records signified a varied range of pollen richness (range between 2-11 per species) and abundance (average of 400 pollen grains per species) between butterfly species as well as the date they were captured.

### Interaction Network and Statistical Analysis

We constructed a quantitative bipartite network using 19 pollinator (butterfly) and 29 plant taxa, highlighting interaction frequency, where the thickness of the bars represents pollen species abundance and linkage indicates the frequency of interaction (Fig. 2). Some relevant parameters examined in the bipartite interaction network include connectance, nestedness, and degree of specialization (Appendix 1 Table A2). In addition, to explore how plant-pollinator interactions changed over time, specimens captured before 2000 and after 2000 were analyzed separately and network parameters were calculated for the respective networks (Fig. 3). Connectance values increased from 0.20 to 0.23 between common species that were captured in both time intervals. High H2’ values for both time periods indicate substantially specialized networks (0.68). The nestedness value also increased from 22.82 to 27.37. Pollen richness over time was also evaluated for the 19 butterfly species, and for most species there was in increase in the number of pollen species over time.

### Focused comparison of field observations to pollen analysis

The composition of the types and abundances of pollen found on the two butterfly species involved in the field observations differed substantially from the plants these two species were observed interacting with. For *Lycaeides anna*, a total of 145 pollen grains representing five plant taxa were found on the collected specimens. This is in contrast to the field observations where individuals were found interacting with three different flower species, though two of the plant taxa from the different study types overlapped, *Solidago californica* and *Aster sp*. (Asteraceae). For *L. melissa*, a total of 119 pollen grains were counted and a total of seven plant species were found on the collected specimens. Though many of the individuals were encountered while flying or on other plants during the observation period for this species, they were only observed interacting with the flowers of *Medicago sativa*.

## Discussion

Plant-pollinator interactions are some of the most important interactions for the maintenance of terrestrial biodiversity. Constructing accurate pollination networks across variable habitats provides important estimates of interactions that can help understand pollinator responses to habitat reduction or fragmentation, climate change, invasions by exotic species, and changing communities generally. We demonstrate that collecting pollen from museum specimens can be a novel and effective means for characterizing and studying pollination interactions and niche breadth in light of changing environmental pressures over time.

Our primary objectives were to summarize the observed pollen-butterfly interaction networks and to determine changes in frequency and richness of those interactions over the last 100 years in California and Nevada. We found a substantial change in pollen richness and abundance on museum specimens collected in different decades. We estimated similar pollen richness over time for a number of different butterfly species, including *Cercyonis pegala*, *Nathalis iole*, *Junonia coenia*, *Nymphalis antiopa*, *Vanessa carye*, *Lycaena helloides*, *Speyeria coronis* and *Oeneis chryxus* from 1910 to present, which suggested that niche breadth and pollen interactions for these species have changed very little in these species over the last 100 years. However, a large increase in pollen richness was observed for several species, including *Hesperia juba*, *Speyeria egleis* and *Cercyonis sthenele*, indicating some pollinator species may be expanding their niche breadth in response to, or rather in spite of, environmental change (Table S2). Alternatively, plant communities have shifted such that available pollen richness has increased and is reflected on the specimens gleaned from butterflies.

Although our results indicate some mild increases in network specialization over time, most butterflies including specialized species may utilize novel plants when focal resources are scarce, and such switching may help stabilize ecosystem processes (Dunne et al., 2002; Tylianakis et al., 2010). The networks observed in our study had connectance values around 0.20, which is higher than what is commonly reported in the literature for visitation networks (Dunne et al., 2002). This may indicate that pollen-butterfly interactions are comprised of a higher degree of interaction complexity that what is estimated from visitation networks. We also found that the networks analyzed had a high level of nestedness, which is linked to high levels functional redundancy, which is also a stabilizing factor for biotic communities.

Another objective of this study was to investigate how climate change has impacted pollen-butterfly interactions over time. In 2015, the snowpack in the Sierra Nevada mountains was only five percent of its historical average, a clear indication of long-term drought in the region (Belmecheri et al., 2016). This is part of an overall trend of increasing drought in the Sierra Nevada and Great Basin regions. Droughts are known to negatively affect pollinators by decreasing the quantity and quality of floral rewards (Phillips et al., 2018). Our study documented clear shifts in pollen-butterfly interactions before and after the onset of drought, based on the more common specimens of Lepidoptera at the UNRMNH (Fig.3). We found that the interaction network after 2000 is characterized by decreased pollen loads compared to before 2000, except in a few butterfly species. We even observed some specimens that were captured after 2000 with no pollen on their body. Pollen-butterfly interaction links decreased in *Polygonia satyrus, Phyciodes mylitta, Nymphalis antiopa,* and *Cercyonis sthenele*. For two species, *Hesperia juba* and *Speyeria egleis*, pollen richness and abundance increased.

Our networks indicated that species that were captured after 2000 are more specialized, which could be a consequence of lower resource availability. The long-term increase in pollen richness with low abundance is correlated with increased drought in California and Nevada in the last two decades. These areas are facing drought that has resulted in increasing and more intense wildfires, and warming temperatures that cause precipitation to fall as rain rather than snow. These changing climate factors are strongly impacting plant communities with which pollinators interact, often priming them for invasion by exotic species and a decrease in abundance of native plants. While butterflies may be confronting a narrower nectar and pollen niche breadth, some may form novel interactions with new plant resources, increasing numbers of interactions over time; such increases in interactions are often associated with increasing stability. Network connectance, which depends on potential interactions in the network, also increased after 2000, and was partly driven by increased niche breadths of a few species, like *S. egleis*, *H. juba* and *N. antiopa* in the last decades. Related to these patterns, the most common pollen species represented in our networks, *E. nauseosa* (Rubber rabbitbrush) and *Monardella* sp sharply declined in abundance in the last two decades compared to the years before 2000; this may be linked to habitat loss and ecosystem alteration in the last two decades.

Finally, the focused observations of two butterfly species, *L. Melissa* and *L. anna*, along with decades of natural history observations, provided a snapshot of the differences between interactions observed in the field (which were designed to mirror widely used visitation network methods) and interactions inferred from pollen found on the specimens’ bodies. Though there was overlap between the observations and the pollen composition found on *L. anna*, there was no pollen from the larval hostplant (*Medicago sativa*) found on the bodies of *L. Melissa* collected in the field in spite of frequent observed interactions between adult moths and flowers for these species. This result highlights potential shortcomings of visitation networks as well as the method used in this paper; the visitation networks clearly underestimate diversity of pollen picked up by butterflies, and in the case of our method, we did not see pollen abundances associated with frequent visitation of a focal flower. These observations were very limited in temporal and spatial extent, but they underscore the importance of combining multiple methods for estimating true networks.

## Conclusion

In a time of growing concern about insect declines, we present a case for the use of museum specimens in describing changes in interaction networks over time. Much media coverage has been directed at the declines in pollinators in recent decades (Biesmeijer et al., 2006; Colla & Packer., 2008; Wood et al., 2019), and long-term data sets have been used to validate these trends (Cola et al., 2012; Bartomeus et al., 2013; Wood et al., 2019). An advantage of using museum specimens is that the collection events may span decades or even more than a century, allowing for reconstruction of past interaction networks. The use of museum specimens also comes with its disadvantages; pollen may fall off or be removed from specimens, there is a lack of standardization in the collection methods, and a host of other factors that can vary across taxa and collections. Also, there will likely be uneven sampling over the years that could generate data that is highly variable over time. However, every approach to describing and quantifying mutualistic networks has disadvantages, and perfect characterizations of species interaction networks are impossible at broad spatial and temporal scales. This approach combined with the more popular visitation networks, and studies of pollinator effectiveness (Ballantyne et al., 2015), could lead to a thorough understanding of pollination interaction networks and how they change across disturbance gradients.

## Acknowledgments

We are grateful to those who captured and collected butterflies for the museum and to M. Forister for helpful comments on this manuscript. Also, we would like to say thank you to the University of Nevada Reno Museum of Natural History (UNRMNH) for funding this research and giving us this opportunity to work on these valuable collections.

## Data Availability Statement

The data that support the findings of this study are openly available at http://dx.doi.org/10.6084/m9.figshare.13236989 and in the supplementary material of this article.

## Biosketch

**Behnaz Balmaki** is a post-doctoral researcher interested in environmental data analyst with interests in research questions relating to palynology, paleoecology, and paleoclimatology. Her current research is largely focused on pollen–pollinator interaction and pollinator responses to the past and future climate change.

**Lee A. Dyer** uses observational, experimental, and quantitative approaches to study chemically mediated plant-insect interactions in temperate and tropical ecosystems. He enjoys poetry, rock climbing, natural history, and applied statistics.

**Tara Christensen** is a PhD student interested in plant-insect interactions and insect disease dynamics. She is interested in direct and indirect interactions between insect functional guilds and the plants that mediate those interactions.

